# Transcriptome sample statistics of the responses the sugar beet root maggot, *Tetanops myopaeformis* has while experiencing susceptible and resistant reactions with sugar beet, *Beta vulgaris* ssp. vulgaris

**DOI:** 10.1101/2024.11.19.624160

**Authors:** Sudha Acharya, Nadim W. Alkharouf, Muhammad Massub Tehseen, Chenggen Chu, Vincent P. Klink

**Affiliations:** Department of Computer and Information Sciences, Towson University, Towson, MD, 21252, USA; USDA-ARS-NA, Northern Great Plains Research Laboratory, 1307 N 18TH ST Northern Crop Science Laboratory, Fargo, ND 58102, USA; Department of Plant Sciences, North Dakota State University, Fargo, ND 58102, USA; USDA-ARS-NEA-BARC, Molecular Plant Pathology Laboratory, Building 004, Room 122, BARC-West, 10300 Baltimore Ave., Beltsville, MD 20705, USA

**Keywords:** sugar beet root maggot, *Tetanops myopaeformis*, sugar beet, *Beta vulgaris*, transcriptome, resistant, susceptible, RNA-seq

## Abstract

The sugar beet root maggot (SBRM), *Tetanops myopaeformis* (von Röder), is a devastating insect pathogen of sugar beet (SB), Beta vulgaris ssp, vulgaris (*B. vulgaris*), one of only two plants in the world from which significant global raw sugar is produced, $1 billion, U.S., $4.6 B, globally. Experiments reveal the SBRM larval transcriptome experiencing two different susceptible or resistant responses by sugar beet SBRM larvae were sampled at time = 0 hours post infection [hpi]), prior to being introduced to *B. vulgaris* and after infection on F1016 and F1024 (resistant), and F1010 and L19 (susceptible) for 24, 48, and 72 hpi when the larvae were removed for transcriptomic analysis. The transcriptomic analyses included determining the number of reads per sample, mapping the transcripts to the recently sequenced SBRM TmSBRM_v1.0 draft genome, identifying genes that relate to the resistant and susceptible responses. Moreover, the RNA-seq experiments provide data for generating differential expression analyses between the various sample types, thus, yielding an understanding SBRM biology, the development of new control strategies for this pathogen, relationship to model genetic organisms like *Drosophila melanogaster*, relationship to pathogenic non-model organisms, and aid in agronomic improvement of sugar beet for stakeholders.

## Introduction

Beta vulgaris ssp, vulgaris (*B. vulgaris*), sugar beet (SB), Order Carophyllales, Family Amaranthaceae, is one of only two plants, globally, from which sugar is widely produced with a worldwide value of $4.6 B (1). In the U.S., the economically important, but non-native, SB has a value of $1 B, harvested from 1.14 million acres of land. Upon introduction to the U.S., SB was encountered by the native insect pathogen *T. myopaeformis* (SBRM) on which it can complete its life cycle and while it can complete its life cycle on other non-native plant species, the native SBRM host has not yet been identified (2-6). SBRM is the most devastating SB pathogen in North America where it can decrease yield by up to 100%, locally, and of further concern is its increasing geographic spread (7-10). Morphological, anatomical, and phenological details of the SBRM life cycle are available and important for control, management, and eradication. However, genomic information that could be used in biological control measures have not been developed and/or employed on the commercial scale (11-19). Problems in the control of SBRM are also compounded by public opinion pressures, insect resistance to insecticides, and banned insecticide use (10, 20-21). The agricultural control of SBRM is also limited by a scarcity of genetic resistance in SB, although progress is being made (1, 22-25). Furthermore, SB and SBRM annotated genomes provide basic knowledge that is crucial for SB improvement and food security (26-31).

The SBRM (Order Diptera, Family Ulidiidae), belongs to a genus of 15 Palaearctic and Nearctic species with some having pathogenic life cycles (32-36). The shared disease-causing habit of some species of the *Tetanops* genus indicates that the pathogenic life cycle may have aspects that are conserved between its species and are under genetic control which appears to be indicated in a recent published genome sequence and its annotation (30-31). Consequently, it may be possible to understand the pathogenic nature of SBRM in ways that would be facilitated by transcriptomic information. Such inferences have been made in other insects, including model genomic systems like *Drosophila melanogaster* and its related *D. suzukii*, and adapted to non-model systems (37-41).

In the analysis presented here, RNA has been isolated from SBRM larva prior to being introduced to either of 2 different SBRM -susceptible (F1010 and L19) or -resistant (F1016 and F1024) SB genotypes. The SBRM were then allowed to infect SB for 24, 48, or 72 hours post infection (hpi) at which times they were collected for RNA isolation and RNA sequencing. The processed data is presented here, relating to the RNA sequence read counts for each sample type. The RNA-seq analysis of SBRM, as it is experiencing either an incompatible (*B. vulgaris*-resistant) or compatible (*B. vulgaris*-susceptible) reaction to any defence response by SB will provide critical data that researchers can use to understand the SBRM biology under these circumstances in ways originally sought in model organisms (37, 42).

## Materials and methods

### Plant infection

SBRM larvae were collected in mid-June 2022 from a field location close to St. Thomas, ND. After cleaning all larvae using 1% Clorox Germicidal Bleach, the 1- and 2-instar larvae were used for root infestation of *B. vulgaris* F1016 (PI 608437) and F1024 (PI 658654) that are resistant, and F1010 (PI 535818) and L19 (PI 590690) that are susceptible genotypes (43-47). The infestation experiment included three replications for each genotype with three plants infested in each replication. For preparing roots for infestation, seeds were germinated using 1% hydrogen peroxide solution (48), and germinated seeds were planted in a greenhouse room under 16:8 (day:night) light regime with temperature range between 20 – 30°C. Roots were collected 4 weeks after planting. After being cleaned to remove the soil, three roots of each genotype as one replication were placed on a 15 cm × 10 cm, 0.8% agar plate (49). Subsequently, fifteen 1- or 2-instar larvae were added to each plate with 5 larvae per root. All plates were then kept in dark at 28°C. Root and insect samples were collected at 0 hpi (right before infestation), and subsequently at 24, 48, and 72 hpi. All samples were immediately flash frozen into liquid nitrogen and then stored at -80°C before RNA isolation and subsequent RNA-seq data generation.

### RNA isolation

Flash-frozen SBRM larval samples were sent to Omega Bioservices Inc., 400 Pinnacle Way, Ste 425, Norcross, GA 30071 for RNA isolation, quality assurance, and RNA sequencing according to Alsherhi et al. (2019). In brief, the RNA isolation implemented a well-established protocol for RNA isolation and library preparation to achieve high-quality sequencing data. The Omega Biotek E.Z.N.A.^®^ Total RNA Kit (Omega Bio-tek) was used to extract total RNA from the samples, following the manufacturer’s protocol. The concentration and integrity of the RNA were assessed using a Nanodrop 2000c spectrophotometer (Thermo Scientific Inc.) and an Agilent 4150 TapeStation instrument (Agilent Technologies), respectively.

### RNA library preparation

For library generation, up to 1 mg of total RNA was used according to the manufacturer’s instructions for the NEBNext^®^ Poly(A) mRNA Magnetic Isolation Module E7490L and NEBNext^®^ Ultra™ II Directional RNA Library Prep Kit for Illumina^®^ E7760L (New England Biolabs Inc.). Quality and quantity evaluation of the libraries were conducted using the High Sensitivity D1000 Screen Tape on an Agilent 4150 TapeStation instrument. Subsequently, the libraries underwent normalization, pooling, and were sequenced with Illumina Novaseq X Plus instrument (Illumina, Inc.) following the manufacturer’s recommendations.

### RNA-seq data processing

For the study presented here, the RNA-seq data analysis process used Geneious prime (https://www.geneious.com/), version 2024.0 with the steps of that pipeline detailed at https://www.geneious.com/series/expression-analysis. The analysis process presented here involved sequence trimming, alignment, and counting. Trimming was used to increase the read’s mapping rate by eliminating adapter sequences and removing poor-quality nucleotides. The alignment was performed to the SBRM TmSBRM_v1.0 draft genome. After mapping the reads, they were assigned to a gene or transcript in a process known as counting or quantification. This step was followed by a normalization procedure employed to remove possible sequencing bias.

## Results

### RNA-seq data processing

The experimental pipeline is presented (**Figure 1**). The RNA-seq analysis has resulted in acquiring data for each of the 39 samples (**Table 1**). The reads have then been mapped to the recently sequenced SBRM TmSBRM_v1.0 draft genome. This analysis has allowed for the generation of a general assessment of gene activity on the SBRM TmSBRM_v1.0 draft genome, aided by its annotation.

**Table 1.** Transcriptome statistics.

| Sample NO. | SBRM use | SB genotype | Outcome | Time point | Total processed reads | Assembled (Used reads) | % Assembled | Unassembled | % Unassembled |
| --- | --- | --- | --- | --- | --- | --- | --- | --- | --- |
| 1 | SBRM control | no SB* | n/a | 0 hpi | 41,366,384 | 29,614,682 | 71.59117896 | 11,751,702 | 28.40882104 |
| 2 | SBRM control | no SB* | n/a | 0 hpi | 45,730,450 | 32,709,379 | 71.52647525 | 13,021,071 | 28.47352475 |
| 3 | SBRM control | no SB* | n/a | 0 hpi | 38,753,604 | 27,516,689 | 71.00420647 | 11,236,915 | 28.99579353 |
| 4 | SBRM infested | F1024 | resistant | 24 hpi | 44,942,632 | 32,898,251 | 73.20054375 | 12,044,381 | 26.79945625 |
| 5 | SBRM infested | F1024 | resistant | 24 hpi | 43,740,534 | 31,705,744 | 72.48595548 | 12,034,790 | 27.51404452 |
| 6 | SBRM infested | F1024 | resistant | 24 hpi | 41,578,980 | 29,385,047 | 70.67284238 | 12,193,933 | 29.32715762 |
| 7 | SBRM infested | F1016 | resistant | 24 hpi | 42,906,878 | 30,892,119 | 71.99805821 | 12,014,759 | 28.00194179 |
| 8 | SBRM infested | F1016 | resistant | 24 hpi | 45,633,030 | 33,527,362 | 73.47169802 | 12,105,668 | 26.52830198 |
| 9 | SBRM infested | F1016 | resistant | 24 hpi | 37,947,020 | 27,188,332 | 71.64813469 | 10,758,688 | 28.35186531 |
| 10 | SBRM infested | F1010 | susceptible | 24 hpi | 45,366,574 | 32,685,868 | 72.04834996 | 12,680,706 | 27.95165004 |
| 11 | SBRM infested | F1010 | susceptible | 24 hpi | 39,166,906 | 27,856,471 | 71.12247008 | 11,310,435 | 28.87752992 |
| 12 | SBRM infested | F1010 | susceptible | 24 hpi | 39,703,208 | 28,549,398 | 71.90703079 | 11,153,810 | 28.09296921 |
| 13 | SBRM infested | L19 | susceptible | 24 hpi | 38,527,460 | 27,743,031 | 72.00846098 | 10,784,429 | 27.99153902 |
| 14 | SBRM infested | L19 | susceptible | 24 hpi | 39,363,640 | 28,339,559 | 71.99425409 | 11,024,081 | 28.00574591 |
| 15 | SBRM infested | L19 | susceptible | 24 hpi | 46,319,624 | 33,639,849 | 72.62547943 | 12,679,775 | 27.37452057 |
| 16 | SBRM infested | F1024 | resistant | 48 hpi | 41,549,788 | 28,818,653 | 69.35932621 | 12,731,135 | 30.64067379 |
| 17 | SBRM infested | F1024 | resistant | 48 hpi | 40,742,024 | 28,661,525 | 70.34880005 | 12,080,499 | 29.65119995 |
| 18 | SBRM infested | F1024 | resistant | 48 hpi | 45,219,784 | 32,076,880 | 70.93550027 | 13,142,904 | 29.06449973 |
| 19 | SBRM infested | F1016 | resistant | 48 hpi | 38,964,178 | 27,309,685 | 70.08921117 | 11,654,493 | 29.91078883 |
| 20 | SBRM infested | F1016 | resistant | 48 hpi | 40,442,336 | 29,203,809 | 72.21098455 | 11,238,527 | 27.78901545 |
| 21 | SBRM infested | F1016 | resistant | 48 hpi | 39,566,764 | 29,027,903 | 73.36435954 | 10,538,861 | 26.63564046 |
| 22 | SBRM infested | F1010 | susceptible | 48 hpi | 42,317,332 | 30,013,792 | 70.92552999 | 12,303,540 | 29.07447001 |
| 23 | SBRM infested | F1010 | susceptible | 48 hpi | 43,407,312 | 30,678,669 | 70.67626993 | 12,728,643 | 29.32373007 |
| 24 | SBRM infested | F1010 | susceptible | 48 hpi | 42,600,156 | 29,429,372 | 69.08277988 | 13,170,784 | 30.91722012 |
| 25 | SBRM infested | L19 | susceptible | 48 hpi | 41,129,322 | 29,218,917 | 71.04157224 | 11,910,405 | 28.95842776 |
| 26 | SBRM infested | L19 | susceptible | 48 hpi | 38,589,096 | 27,004,810 | 69.98041623 | 11,584,286 | 30.01958377 |
| 27 | SBRM infested | L19 | susceptible | 48 hpi | 41,630,316 | 29,920,707 | 71.87239943 | 11,709,609 | 28.12760057 |
| 28 | SBRM infested | F1024 | resistant | 72 hpi | 43,111,724 | 29,379,202 | 68.1466647 | 13,732,522 | 31.8533353 |
| 29 | SBRM infested | F1024 | resistant | 72 hpi | 41,201,980 | 28,859,219 | 70.0432819 | 12,342,761 | 29.9567181 |
| 30 | SBRM infested | F1024 | resistant | 72 hpi | 39,162,290 | 27,308,993 | 69.73288079 | 11,853,297 | 30.26711921 |
| 31 | SBRM infested | F1016 | resistant | 72 hpi | 40,314,630 | 28,519,330 | 70.741887 | 11,795,300 | 29.258113 |
| 32 | SBRM infested | F1016 | resistant | 72 hpi | 42,578,254 | 29,420,002 | 69.09630912 | 13,158,252 | 30.90369088 |
| 33 | SBRM infested | F1016 | resistant | 72 hpi | 42,440,440 | 28,561,396 | 67.29759635 | 13,879,044 | 32.70240365 |
| 34 | SBRM infested | F1010 | susceptible | 72 hpi | 44,288,560 | 29,460,791 | 66.52009232 | 14,827,769 | 33.47990768 |
| 35 | SBRM infested | F1010 | susceptible | 72 hpi | 44,569,173 | 29,849,449 | 66.97330686 | 14,719,724 | 33.02669314 |
| 36 | SBRM infested | F1010 | susceptible | 72 hpi | 39,120,669 | 25,883,749 | 66.16387107 | 13,236,920 | 33.83612893 |
| 37 | SBRM infested | L19 | susceptible | 72 hpi | 40,681,422 | 28,238,881 | 69.41468516 | 12,442,541 | 30.58531484 |
| 38 | SBRM infested | L19 | susceptible | 72 hpi | 41,919,080 | 28,609,391 | 68.24909087 | 13,309,689 | 31.75090913 |
| 39 | SBRM infested | L19 | susceptible | 72 hpi | 38,559,366 | 27,700,699 | 71.83909352 | 10,858,667 | 28.16090648 |

\*The SBRM larvae were not exposed to sugar beet.

**Figure 1.**
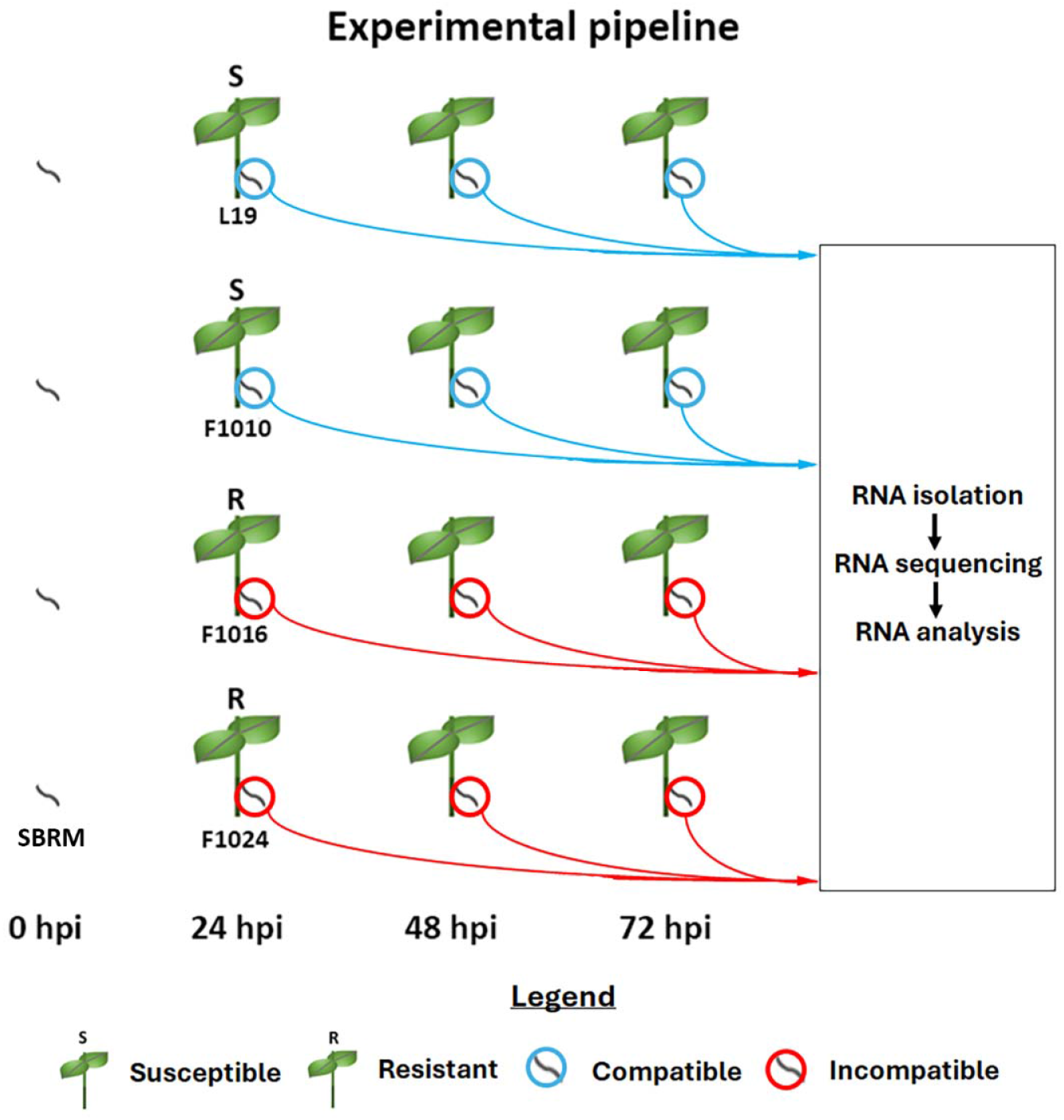
Experimental pipeline. The SBRM-susceptible L19 and F1010, and SBRM-resistant F1016, and F1024 *B. vulgaris* genotypes are shown. The respective compatible and incompatible SBRM are encircled by a blue or red ring. At t = 0 hpi, the SBRM were collected before any introduction to *B. vulgaris*. Thus, the SBRM are shown to not be closely associated with *B. vulgaris*. SBRM are subsequently shown to be in direct contact with *B. vulgaris* at the t = 24, 48, and 72 hpi time points. The samples were collected for transcriptomic study that involved RNA isolation, sequencing, and analysis.

## Discussion

RNA-seq data has been obtained from each of the 39 examined samples. The analysis has identified a range in total processed reads per sample of 37,947,020 to 46,319,624 in total assembled (used) reads per sample of 25,883,749 (66.16%) to 29,385,047 (70.67%), and a range in total unassembled reads per sample of 10,538,861 (26.64%) to 12,303,540 (29.07%). The range in average used reads per time point was 23,246 (L19 resistant, 24 hpi) to 24,525 (F1010 resistant, 72 hpi), 5.22%. Therefore, the sample read quantity is similar between the different samples. From these data, further processing is possible, allowing for an idea of differential expression of genes during the susceptible and resistant reactions. The analyses will allow for the identification of genes, gene pathways, and biological processes which may or may not fall under gene pathways to be identified. The identification of the genes, gene pathways, and biological processes will allow scientists to devise management, control, and biological assays for SBRM much in the same way that has been done for other devastating agricultural pathogens (51-53).

## Ethics statement

The authors have read and follow the ethical requirements for publication in Bioinformation and confirming that the current work does not involve human subjects, animal experiments, or any data collected from social media platforms.

## Author credit statement

**SA** Methodology; Software; Validation; Formal analysis; Investigation; Resources; Data Curation; Writing - Original Draft

**NA** Methodology; Software; Validation; Formal analysis; Investigation; Resources; Data Curation; Writing - Original Draft

**MT** Investigation; Resources

**CC** Investigation; Resources, Supervision; Project administration; Funding acquisition

**VK** Conceptualization; Methodology; Resources; Visualization; Supervision; Project administration; Funding acquisition; Writing - Original Draft

## Acknowledgements

This work was supported by the USDA-ARS NP 8042-21220-262-000D project to VK and USDA-ARS NP 3060-21000-045-000D to CC. The mention of trade names or commercial products in this publication was solely for the purpose of providing specific information and does not imply recommendation or endorsement by the United States Department of Agriculture. USDA is an equal opportunity provider and employer.

